# K-mer based prediction of *Clostridioides difficile* relatedness and ribotypes

**DOI:** 10.1101/2021.05.17.444522

**Authors:** Matthew. P. Moore, Mark H. Wilcox, A. Sarah Walker, David. W. Eyre

## Abstract

Comparative analysis of *Clostridioides difficile* whole-genome sequencing (WGS) data enables fine scaled investigation of transmission and is increasingly becoming part of routine surveillance. However, these analyses are constrained by the computational requirements of the large volumes of data involved. By decomposing WGS reads or assemblies into k-mers and using the dimensionality reduction technique MinHash, it is possible to rapidly approximate genomic distances without alignment. Here we assessed the performance of MinHash, as implemented by sourmash, in predicting single nucleotide differences between genomes (SNPs) and *C. difficile* ribotypes (RTs). For a set of 1,905 diverse *C. difficile* genomes (differing by 0-168,519 SNPs), using sourmash to screen for closely related genomes, at a sensitivity of 100% for pairs ≤10 SNPs, sourmash reduced the number of pairs from 1,813,560 overall to 161,934, i.e., by 91%, with a positive predictive value of 32% to correctly identify pairs ≤10 SNPs (maximum SNP distance 4,144). At a sensitivity of 95%, pairs were reduced by 94% to 108,266 and PPV increased to 45% (maximum SNP distance 1,009). Increasing the MinHash sketch size above 2000 produced minimal performance improvement. We also explored a MinHash similarity-based ribotype prediction method. Genomes with known ribotypes (n=3,937) were split into a training set (2,937) and test set (1,000) randomly. The training set was used to construct a sourmash index against which genomes from the test set were compared. If the closest 5 genomes in the index had the same ribotype this was taken to predict the searched genome’s ribotype. Using our MinHash ribotype index, predicted ribotypes were correct in 780/1000 (78%) genomes, incorrect in 20 (2%), and indeterminant in 200 (20%). Relaxing the classifier to 4/5 closest matches with the same RT improved the correct predictions to 87%. Using MinHash it is possible to subsample *C. difficile* genome k-mer hashes and use them to approximate small genomic differences within minutes, significantly reducing the search space for further analysis.

**Impact statement:** The genetic code, or DNA, of bacteria is increasingly used to track how infection spreads and to guide infection control interventions, as similar or identical DNA sequences are expected in samples from pair of individuals related by transmission. While obtaining the DNA sequence for bacteria is increasingly straightforward, comparing thousands or even millions of sequences requires substantial computing power and time using current approaches. Here we describe how a method for summarising sequencing data, MinHash, can be used to rapidly reduce the number of possible close sequence matches in *Clostridioides difficile*, an important healthcare-associated pathogen. It can also be used to approximate traditional schemes used to classify *C. difficile* into smaller subgroups in transmission analyses, such as ribotyping.

**Data summary:** The authors confirm all supporting data, code and protocols have been provided within the article or through supplementary data files.

## Introduction

Whole genome sequencing (WGS) has transformed our understanding of the epidemiology of many bacterial pathogens. For example, *Clostridioides* (formerly *Clostridium*) *difficile*, previously believed to be transmitted predominantly from other hospitalised cases, has been shown to be mostly acquired from other sources(1). WGS also facilitates outbreak investigations, e.g. of enteric pathogens(2–4) and highly drug-resistant gonorrhoea(5,6), and can support specific infection control interventions(7) and surveillance(8). However, all these potential applications rely on being able to identify closely genetically related infections against an ever-growing collection of previously sequenced genomes. To date most studies have relied on constructing phylogenies on the basis of single nucleotide polymorphisms (SNPs), however such approaches are computationally demanding, and increasingly impractical for large scale comparisons of genomes for timely public health or infection control interventions.

One potential approach to this challenge is to ‘divide and conquer’, such that a rapid screen is used to define which subgroup of a species a genome of interest is from, reducing the number of genomes that need to be considered in a more detailed analysis. For example, multi-locus sequencing typing (MLST), based in *C. difficile* on seven housekeeping genes,(9) can be used to rapidly type isolates from sequencing reads without the need for mapping or assembly(10). MLST can eliminate the possibility of an outbreak if isolate sequences have diverse sequence types (STs) but lacks the discriminatory precision to identify whether sequences of the same ST constitute a likely outbreak. Core genome MLST (cgMLST) expands on MLST providing, in theory, a standardised set of 2000-3000 *C. difficile* core genes(11) for far greater differentiation of isolates by genomic data. The selection of cgMLST loci for schema creation, quality assessment and designation of profiles, and ongoing curation of cgMLST schemes, have been made possible by open-source software(12). However, there are multiple cgMLST schemes for *C. difficile* that are not readily inter-operable with open-source software(12) due to differences in reference genome annotation and gene selection protocols. To address these limitations and improve cgMLST for outbreak detection, a hash based cgMLST scheme has been developed for *C. difficile*, removing the need for a centralised, curated database(13). Other typing schemes can also be used to subdivide isolates. For *C. difficile*, PCR ribotyping(14) remains the standard for non-WGS-based surveillance. However, identifying PCR ribotypes directly from WGS based on short reads is challenging. An *in silico* ribotyping method has been developed to detect ribotype (RT) 027, but not ribotypes more generally(15). There is not a perfect 1:1 correspondence between ribotypes and STs, but common equivalents, e.g. ST1/RT027 are well described.

We therefore investigated the performance of alternative approaches based on k-mers for screening isolates to identify subsets with more closely related WGS. K-mers are fragments of sequence data of length k. Therefore, these approaches are potentially species agnostic, and could be deployed widely, without the need for species-specific schemes such as MLST, cgMLST or ribotyping. They also have the advantage that they do not necessarily require prior genome assembly or alignment as required for cgMLST or SNP based analyses respectively. Bacterial genome sequences contain millions of k-mers and so dimensionality reduction is used to improve performance. The MinHash dimensionality reduction method is a form of local sensitivity hashing(16) that was first applied to comparative genomics with the development of MinHash extension, Mash(17). With Mash, and similar k-mer-based dimensionality reduction techniques developed subsequently(18–20), it is now possible to quickly compare tens of thousands of genomes and rapidly identify organisms in metagenomic sequencing projects without alignment(21). These methods depend on a genome k-mer ‘sketch’ (a random subset of all k-mers of a certain size) and as such precision, in theory, may be increased by increasing the sketch size with the cost of increased compute time. The precision of these methods also depends upon biological factors such as how correlated core genome SNP distances are with accessory gene differences within a species or lineage of interest(22).

Here we test the performance of k-mer based algorithms in *C. difficile* and assess whether these algorithms can be used to approximate the most common typing scheme in use, ribotyping, and identify sequence pairs within SNP thresholds in order to provide backwards compatibility with existing surveillance systems and identify groups of closely related isolates for in depth phylogenetic characterisation.

## Methods

### Data

Previously sequenced genomes from studies of clinical isolates in the CD-link study (six UK hospitals) (n=973), Leeds (n=1,876), *C. difficile* UK ribotyping network (n=684), EUCLID and CLOSER studies (n=678), Leeds and ECDC collections (n=369), RT078 genomes from the NCBI short read archive (n=68) and RT244 from Australia (n=16) with available ribotypes were considered for inclusion (n=4,655). The large majority of ribotypes in the study were assigned by the national *C. difficile* reference laboratory in Leeds, UK using capillary gel electrophoresis as previously described(23). Potential laboratory errors were identified by comparing *in silico* MLST results with known ribotypes. Details of included sequences, short-read archive accession numbers and ribotypes are provided in Table S1.

### Bioinformatic processing

K-mer based comparisons were undertaken with and without prior genome assembly. Use of assemblies potentially allows most sequencing errors to be removed first, but is more computationally intensive. Sequencing adapters were removed using BBDuk(24). Reads were assembled using Velvet(25) with Velvet optimiser (https://github.com/tseemann/VelvetOptimiser), without prior read quality trimming or filtering. Contigs <1000bp in length were filtered out of assemblies to avoid fragmented assemblies. Assembly quality was assessed using QUAST-5.0.2(26). Assemblies <3.7Mb or >4.7Mbp total assembly size were excluded to avoid genomes with insufficient data or potential contamination (n=163 and n=53 respectively). Assemblies with >1000 contigs were also excluded to avoid excessively fragmented assemblies (n=109). Multi-locus sequence types (STs) were determined *in silico* from assemblies using mlst(27) and the scheme of Griffiths *et al*.(9). Those without a determinable *C. difficile* MLST were excluded (n=297). We screened the dataset for potential sample mislabelling by looking for incongruent ST-ribotype pairs with a novel ST-RT network approach. The Python package NetworkX(28) was used to generate a weighted, undirected network with each ST and RT as a node in the network. Each genome sharing an ST and RT added an edge and an edge weight of one to the ST and RT nodes. Potentially mislabelled samples were screened for as edges with low weights where there was strong evidence for an alternative ST-RT relationship: for all connected components with >3 nodes, an edge (connecting node A and B) was considered potentially spurious if the edge weight was less than the sum of edge weights connected to node A divided by ten and also less than the sum of edge weights connected to node B divided by ten. Samples were excluded on this basis (n=96), as re-running the sequencing and ribotyping for this current study was not practical. A random sample of 1905 genomes were taken to assess k-mer vs SNP distances. For analyses without prior genome assembly reads were quality trimmed using TrimGalore!(29,30) with minimum Phred Score 30 and low abundance k-mers removed using khmer trim low abundance(31,32) requiring 10 or more copies of each k-mer.

### Comparison of K-mer based distances with SNP distances

We assessed the performance of several k-mer based approaches. Sourmash(20) was used to generate MinHash Jaccard distances.

To explore explanations for discrepancies between k-mer based metrics and SNPs we simulated perfect sequence reads and reads with sequencing errors using wgsim(33). Three sets of 500 *C. difficile* simulated genome read datasets were simulated using the reference genome 630(34) (length = 4,290,252bp) with a mutation rate set randomly to achieve an expected number of SNPs of between 1 and 500 from the reference genome. Coverage depth was set to 70x. The first simulations comprised SNPs only, the second SNPs with 15% of polymorphisms as indels and the third SNPs with a sequencing error rate of 0.01. In simulations with indels, the default probability that an indel is extended (0.30) was applied. Simulated assemblies were generated by addition of simulated variants to the CD630 genome sequence from wgsim output for each genome. Real pairwise SNP and indel differences between simulated reads and assemblies were recorded from wgsim output.

We analysed the capacity of the k-mer based algorithms to identify sequence pairs within SNP thresholds such as ≤10 SNPs, i.e. samples sufficiently closely related to be related by transmission within the last 5 years(1). Results were also generated for ≤50 SNPs and ≤100 SNPs for comparison. For varying thresholds in the sourmash distance (denoted d^kmer^), we evaluated the number of true positive pairs (TP), i.e. ≤10 SNPs and distance ≥ d^kmer^ (d^kmer^=1 denotes smallest sourmash distance possible and d^kmer^<1 increasingly dissimilar to d^kmer^=0) and false positive pairs (FP), i.e. >10 SNPs and distance ≥ d^kmer^. Sensitivity, specificity and positive predictive value (PPV) were calculated. The PPV value represents the proportion of genome pairs ≥ d^kmer^ that are true positives. The reduction in search space was calculated as the proportion of overall genome pairs < d^kmer^. NetworkX(28) was used to cluster genomes by sourmash distance. Full k-mer hash set comparisons were performed with Dashing(18) khash for comparison. A large sourmhash k-mer size was selected (k = 51) for sensitivity and the optimal sketch size estimated as the smallest sketch size (s = 2000) approximating the performance of full k-mer hash sets.

Unique k-mers in a genome can arise from SNPs and indels, but also from other genetic material (e.g. accessory genes, contamination or insertions). We evaluated the size of the association between greater than expected k-mer distances versus SNPs and the accessory genes present. Accessory genes were predicted from assemblies using roary(35) with the following settings: core genes were defined as those gene clusters present in ≥99% isolates; the blast percent identity to designate genes as the same was set to ≥95% and was repeated at 75% and 50% blast identity. We also evaluated SNP and k-mer distances between sequence data generated from repeated sequencing of the same DNA or from repeated sequencing of the same isolate (Table S1).

### Ribotype prediction

We assessed if k-mer based distances could be used to predict ribotypes directly from sequence reads. Genome sketch signatures were generated with sourmash(20) compute (K=51, sketch size = 2000) for all sequenced genomes. An index (sbt) of all genome sketches was generated with sourmash index and searched with sourmash search, with default parameters. Genome assemblies that passed quality filtering and that had reliable ribotypes (n=3937) were randomly divided into training (n=2,937) and test datasets (n=1000). The training dataset was used to generate a sourmash index. We then compared each sequence in the test dataset to all in the training dataset (sourmash index). The RT of the five most closely related genomes (using sourmash search) in the training dataset were identified, considering the accuracy of sourmash search determined ribotypes based on requiring all 5 closest matches to be the same ribotype and requiring only 4/5 matches to be the same.

## Results

### k-mer distances versus SNPs in simulated data

We used simulated data to initially define the best-possible performance expected from k-mer based methods before assessing real-world performance. We simulated 500 genomes without sequencing errors or indels, each containing from 1 to 500 SNPs from the 630 reference genome, yielding 124,750 pairs of genomes between 2 to 1111 SNPs different. We evaluated the ability of k-mer distances to identify the 24 (0.02%) simulated genomes within ≤10 SNPs. Comparing sourmash signatures from assemblies, at the largest mash distance (0.999) with 100% sensitivity to identify all 24 ≤10 SNP pairs, the search space was reduced by 98.5% (to 1841 from 124,750 genome pairs), with a PPV of 1.3% (24/1841) (Table S4, Figure S4). As expected (assuming no SNPs are within k base pairs of one another), the full jaccard distance from all k-mers (dashing khash) predicted ≤10 SNP pairs with 100% sensitivity and a PPV of 100%. In simulated genomes with SNPs and indels, and mash distance threshold 0.999, the search space reduction was 98.4% (1970/124,750), with a PPV of 1.9% (37/1970) (Table S4, Figure S4). Using full k-mer jaccard distance for this simulation at 100% sensitivity predicted ≤10 SNP pairs with a PPV of 56.1% (37/66). Performance for the SNPs only and SNPs and indels simulations were very similar when sourmash signatures were generated directly from sequencing reads (Table S4, Figure S4). A third set introducing high sequencing error (at a rate of 0.01, with x70 coverage) markedly reduced performance from reads by comparison (Table S4, Figure S4). The simulation had only SNPs and a high level of sequencing errors, and genome pairs differed by 3 to 1046 SNPs (24 pairs ≤10 SNPs) (Table S4, Figure S4). Performance at 100% sensitivity (mash distance threshold 0.105), produced only a 10% reduction in search space (111,502/124,750 genome pairs) with a PPV of 0.02% (24/111,502).

### k-mer distances versus SNPs in real data

A random subsample of *C. difficile* genomes (n=1,905) was selected to assess the precision of mash distances at predicting small SNP cut-offs in real data. The genomes ranged from 0 to 168,519 SNPs different from one another; of 1,813,560 pairs, 52,020 (2.9%) were within ≤10 SNPs, in 172 clusters, each containing a median (range) 3 (1-39,716) pairs.

Using sourmash signatures from assembled genomes, the largest mash distance (0.884) with 100% sensitivity for identifying all ≤10 SNP pairs, reduced the search space by 93.9% (to 109,914/1,813,560 genome pairs) with a PPV of 32.1% (52,020/161,934) (Figure 1, Table 1, Table S3). Pairs ≥0.884 sourmash distance apart fell into 49 clusters containing a median (range) 28 (1 - 99,681) pairs. The largest SNP difference within a sourmash-identified cluster was 4,144. At the largest sourmash distance (0.973) with ≥95% sensitivity to identify ≤10 SNP pairs, the search space was reduced by 94.0% (108,266/1,813,560) with a PPV of 45.8% (49,538/108,266). Pairs ≥0.973 sourmash distance apart fell into 119 clusters containing a median (range) 3 (1 - 88,708) pairs, with the largest pairwise SNP difference within cluster 1,009 (Table 1). Signatures were also generated directly from sequencing reads. This resulted in poorer performance compared with sourmash signatures from assembled genomes (Table 1). Performance was improved by removal of low abundance k-mers (Table 1, Table S5, Figure S5).

**Table 1.** Performance of k-mer based genome comparisons identifying pairs with ≤10 SNPs. Contigs <1000bp were removed from assembled genomes before k-mer hash signatures were generated. Results for k-mer hash signatures are presented from sequencing reads with and without removal of low abundance reads. Core gene k-mer hash signatures were generated from multi-fasta files such that overlapping regions did not generate k-mers

| K-mer source | Sensitivity for including pairs $\leq 10$ SNPs used to determine Sourmash threshold | Sourmash distance threshold | Pairs reduced to (% reduction from 1813560) | Positive Predictive Value (true positives / true positives and false positives) | Clusters identified | Median (range) pairs per cluster | Largest SNP difference within cluster |
| --- | --- | --- | --- | --- | --- | --- | --- |
| **Assemblies** | 100% sensitivity for all $\leq 10$ SNPs | 0.884 | 161,934 ($\downarrow 91.1\%$) | 32.1% (52,020/161,934) | 49 | 28 (1-99,681) | 4,144 |
| **Assemblies** | 95% sensitivity for all $\leq 10$ SNPs | 0.973 | 108,266 ($\downarrow 94.0\%$) | 45.8% (49,538/108,266) | 119 | 3 (1-88,708) | 1,009 |
| **Reads exc low abundance k-mers** | 100% sensitivity for all $\leq 10$ SNPs | 0.780 | 177,940 ($\downarrow 90.2\%$) | 29.2% (52,020/177,940) | 38 | 102 (1-99,681) | 7,705 |
| **Reads exc low abundance k-mers** | 95% sensitivity for all $\leq 10$ SNPs | 0.950 | 125,761 ($\downarrow 93.1\%$) | 39.5% (49,528/125,761) | 72 | 5 (1-96,520) | 1,460 |
| **Reads (all k-mers)** | 100% sensitivity for all $\leq 10$ SNPs | 0.071 | 1,492,029 ($\downarrow 17.7\%$) | 3.5% (52,020/1,492,029) | 1 | 1,492,029 | 105,373 |
| **Core genes** | 100% sensitivity for all $\leq 10$ SNPs | 0.978 | 157,114 ($\downarrow 91.3\%$) | 33.1% (52,020/157,114) | 52 | 40.5 (1-99,681) | 3,202 |

**Figure 1.**
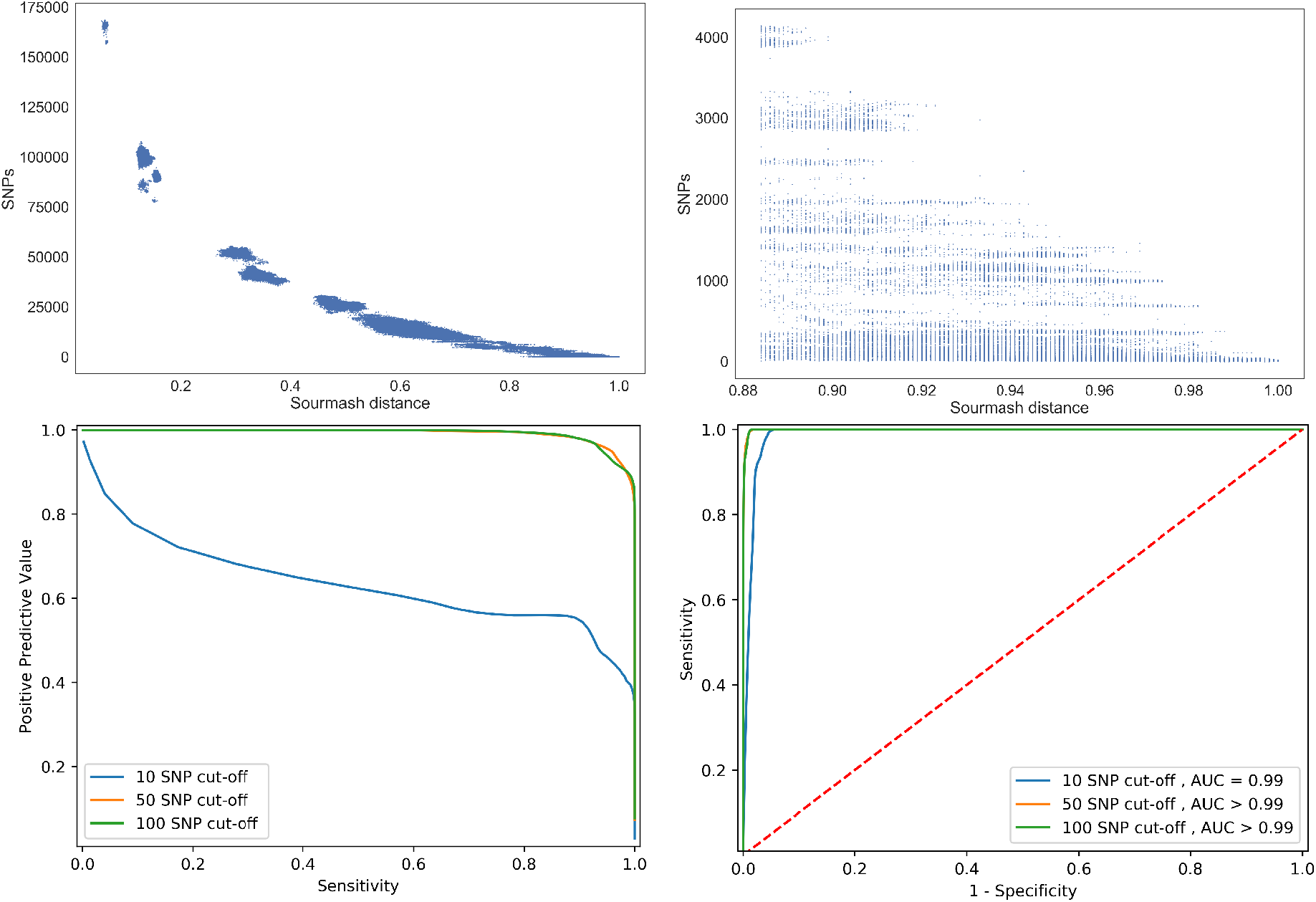
The relationship between and measures of performance of non-redundant pairwise sourmash distances from assembled genomes and their corresponding core genome SNP distances. Top left shows a scatterplot of sourmash distances vs SNP distances (n=1,813,560) comparisons and top right, those ≥0.884 sourmash distance where sensitivity for pairs ≤10 SNPs is 100%. Bottom left shows performance for all sourmash distance thresholds predicting pairs that are ≤100 SNPs, ≤50 SNPs and ≤10 SNPs by positive predictive value vs sensitivity and bottom right the receiver operator curve with area under the curve values rounded to 2 decimal places

To further assess whether k-mer based approaches could improve on existing typing schemes we examined the within-lineage performance of sourmash distance thresholds at 100% sensitivity for predicting ≤10 SNPs between assembled genomes To assess whether performance is being driven by the inclusion of very distant pairs we repeated our analyses within-lineage. This assesses the performance in sets of more similar genome pairs with real data and within-lineage population structure (Figure 2, Table S6). Ribotypes with ≥50 genomes in the sample were included (005, 015, 027, 014, 106, 020, 002, 078, 026 and 001/072; Table 2). For comparison, we considered the random PPV for each lineage assuming that clustering was done on the basis of lineage only: that of no threshold (sourmash distance = 0.000) and estimated the ratio of the PPV achieved by the sourmash threshold vs the random PPV. For the whole dataset (not conditioning on ribotype) the ratio of sourmash:random PPV using a distance with 100% sensitivity to identify pairs with ≤10 SNPs was 11.1 (32.1/2.9). Within ribotypes, the lowest ratio was that of RT078 at 0.8 (24.7/29.4) and the highest RT001/072 at 4.8 (28.9/6) followed by RT014 at 4.1 (12/2.9). In total, in 6/10 ribotypes the sourmash threshold at the 100% sensitivity distance threshold outperformed the random classifier, though none by as much as the whole dataset (Table 2).

**Table 2.** Performance of k-mer based genome comparisons identifying pairs with ≤10 SNPs. All k-mer hash signatures were generated from assembled genomes per ribotype. Random positive predictive value is the performance of Sourmash threshold 0.000, or proportion of the genome pairs with ≤10 SNPs in the dataset

| RT | Median SNPs within lineage | SNP range | Sourmash threshold | Search space reduction | PPV | Random PPV | PPV/Random PPV |
| --- | --- | --- | --- | --- | --- | --- | --- |
| **78** | 17 | 0 - 919 | 0.932 | 6,896/7,503<br>(↓ 8.1%) | 25.0% | 29.4% | 0.8 |
| **5** | 103 | 0 - 5,074 | 0.935 | 940/2,926<br>(↓ 67.9%) | 8.1% | 2.6% | 3.1 |
| **27** | 13 | 0 - 470 | 0.945 | 99,564/99,681<br>(↓ 0.11%) | 40.0% | 39.9% | 1 |
| **26** | 943 | 0 - 6,242 | 0.924 | 464/1,326<br>(↓ 65.0%) | 2.4% | 2.4% | 1 |
| **2** | 65 | 0 - 1,756 | 0.920 | 5,991/8,001<br>(↓ 29.4%) | 9.5% | 6.7% | 1.4 |
| **20** | 44 | 0 - 1,416 | 0.947 | 3,422/3,828<br>(↓ 10.6%) | 8.2% | 7.4% | 1.1 |
| **15** | 4,428 | 0 - 5,700 | 0.884 | 5,061/13,366<br>(↓ 62.1%) | 8.4% | 3.2% | 2.6 |
| **14** | 1,756 | 0 - 4,391 | 0.939 | 1,353/5,671<br>(↓ 76.1%) | 12.0% | 2.9% | 4.1 |
| **106** | 9 | 0 - 1,400 | 0.940 | 14,454/14,878<br>(↓ 2.8%) | 55.0% | 53.2% | 1 |
| **001/072** | 217.5 | 0 - 885 | 0.959 | 740/3,570<br>(↓ 79.3%) | 29.00% | 6.0% | 4.8 |
| **All genomes** | 24,576 | 0 - 168,519 | 0.884 | 161,934/1,813,560<br>(↓ 91.1%) | 32.1% | 2.9 % | 11.1 |

**Figure 2.**
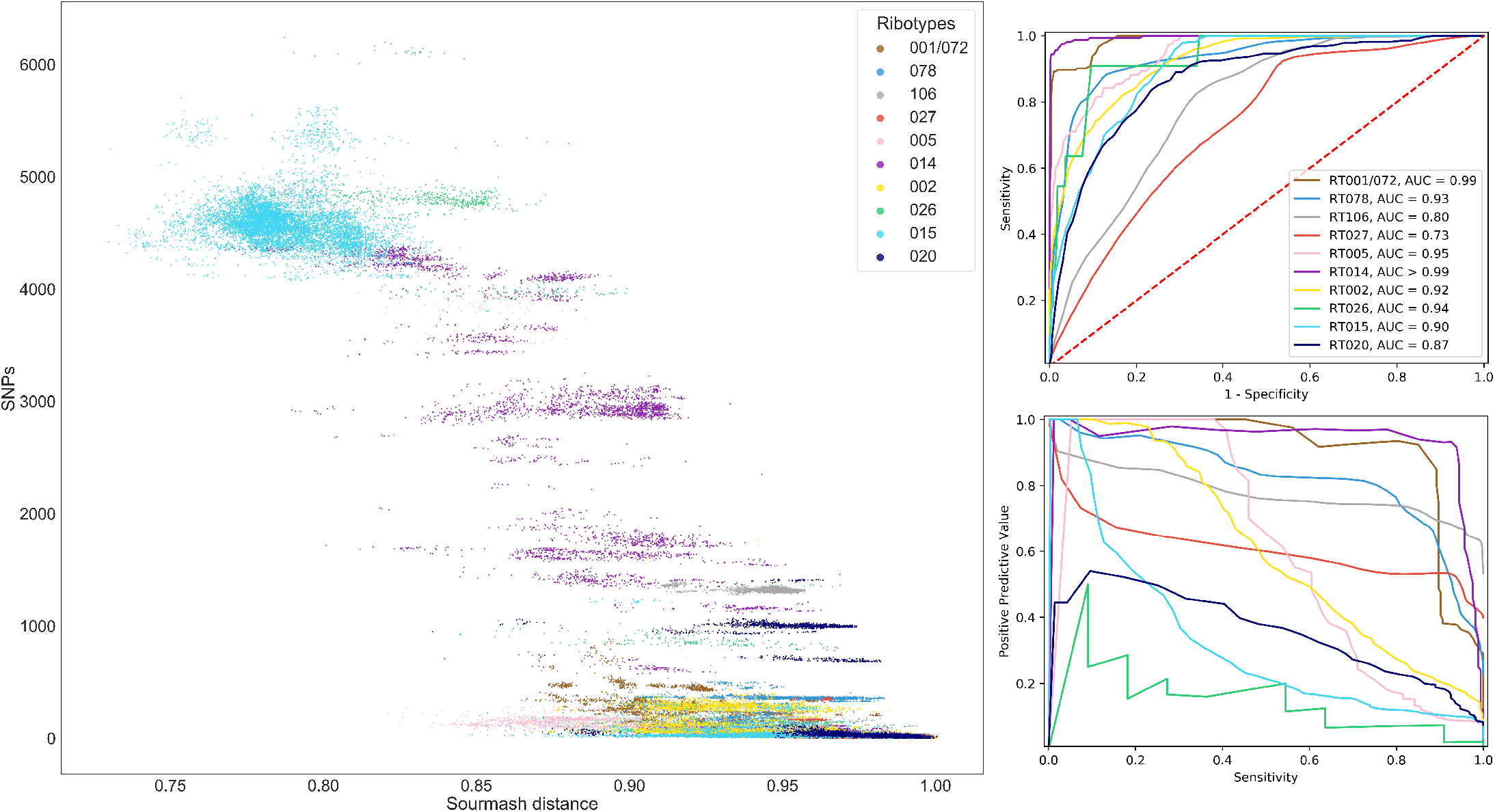
Sourmash and SNP distances within lineage (ribotypes) from assembled genomes for which there were ≥50 genomes per ribotype. Left, a scatterplot shows the relationship between sourmash distance and SNPs within ribotype. Performance of all sourmash thresholds predicting pairs ≤10 SNPs within ribotype is plotted top right with a receiver operator curve and bottom right positive predictive values vs sensitivity. Plots are consistently coloured by ribotype and area under the curve values are rounded to 2 decimal places

One possibility for k-mer based approaches would be to rapidly exclude genome pairs with large SNP distances resulting in clusters of pairs with comparatively smaller distances for further, fine scaled investigations. This process is comparable to clustering by MLST (also rapidly discernible from genome sequences). We compared the SNP distance ranges in sourmash clusters to MLST clusters. The largest SNP distance for instance in sourmash clusters from assembled genomes using a distance with 100% sensitivity to identify pairs with ≤10 SNPs was 4,144 (Table 1). The 161,934 genomes identified at this threshold were clustered by MLST, for those with ≥50 genomes per ST (n = 11). Four of the 11 MLST clusters contained larger SNP distances than the largest within sourmash clusters; the largest in ST3 (13,874) and ST7 (10,296). The remaining 7 ST clusters contained smaller SNP distances than the largest SNP distance within sourmash clusters.

### Impact of gene differences between assemblies on comparisons

We assessed the number of gene differences between genome pairs using roary to assess whether core genome heterogeneity or accessory gene differences were linked to the performance of the sourmash distance approximation. Correlation between sourmash distance dissimilarity was higher with number of SNPs (Spearman’s rho=0.95, p<0.001) than pairwise gene number difference at 95% blast identity (Spearman’s rho=0.79, p<0.001). In pairs within ≤10 SNPs, there was a weaker correlation between sourmash distance dissimilarity and increasing SNPs as expected (Spearman’s rho=0.25, p<0.001) but the correlation between sourmash distance dissimilarity and pairwise gene number difference was higher (Spearman’s rho=0.84, p<0.001). Trends were concordant at 75% and 50% blast identity (Figure S6).

We therefore assessed the performance of k-mer distances using extracted core genome alone. Core genes (gene clusters present in ≥99% isolates) were extracted, signatures were generated based on multi-fasta core gene files and the performance predicting core SNPs assessed. At the largest mash distance (0.978) with 100% sensitivity to identify all ≤10 SNP pairs, the search space was reduced by 91.3% (157,114/1,813,560 genome pairs) with a PPV of 33.1% (52,020/157,114), providing comparable performance with whole genome assemblies overall (PPV 32.1% at 100% sensitivity) (Table 1, Table S7, Figure S7). The correlation between sourmash distance dissimilarity and number of SNPs was also slightly greater (Spearman’s rho=0.97, p<0.001) compared with k-mers generated from assemblies.

We also investigated the number of gene differences reported by roary between replicate sequences. Following quality filtering, 355 assembled genomes were identified from isolates sequenced more than once (median (range) 2 (2-27) times). One genome was selected at random for each isolate and compared to all others from the same isolate (n=259 comparisons). At 95% blast percent identity defining gene presence/absence, the median (range) gene differences between replicate sequences was 95 (28-831), showing the extent to which this metric may be affected by factors other than biological variation. Sourmash distance ranged from 0.914 to 1 with a median of 0.994. Number of genes differing between replicates was highly correlated with sourmash distance (Spearman’s rho=0.85, p<0.001).

### Sourmash Search for *in silico* Ribotype Prediction

As the majority of historical surveillance data has been based on ribotypes it is natural to ask whether sourmash signatures could be used to predict ribotypes from WGS to enable continuity. We investigated whether sourmash search can be used to query a sourmash index (database) of genome signatures with known ribotypes to rapidly assign ribotypes to the search genome signature. All sourmash genome k-mer hash signatures were generated from assembled genomes and sourmash search similarity is reported in estimated percent identity. We evaluated the predicted ribotypes for 1000 randomly selected test genomes from 95 ribotypes by considering the ribotypes of the 5 closest matching genomes in a database (sourmash index) of 2937 genomes from 154 ribotypes. Two rules were tested, requiring 5/5 or 4/5 ribotype concordance among the signatures predicted to be most similar to the search genome signature. The 5 most common test ribotypes searched were RT027 (n=197), RT078 (n=90), RT014 (n=87), RT002 (n=65) and RT015 (n=65). There were 51 test ribotypes with <5 signatures in the database and 46 with <4; both including 18 with none.

Taking the five closest matches per test signature, 4422/5000 (88.4%) matched the searched genome signature, with estimated sourmash percent identity ranging from 77.7-100.0% for correct ribotype matches and 52.3-100.0% for incorrect ribotype matches and an area under the receiver operating curve for sourmash search percent identity of 0.71. Of the 95 test ribotypes, 25 never had a correct individual match resulting in incorrect prediction. In 18 cases this was plausibly due to them not being present in the search database. Of these 25, 23 were present only once in the test dataset and 2 twice.

Requiring all 5 database matches to be the same ribotype resulted in 780 (78%) correctly predicted ribotypes, 200 (20%) undeterminable (not all five top matches the same ribotype) and 20 (2%) incorrect predictions (i.e. all 5 top matches were the same incorrect ribotype) (Table 3). Sensitivity excluding test genomes with ribotypes not present 5 times or more in the search database was 83.3% (780/936). Relaxing the classifier to allow 4/5 concordant matches improved correctly predicted ribotypes to 872 (87.2%) and reduced unknowns to 78 (7.8%), but increased incorrect predictions to 50 (5%) (Table 3). For individual matches, none correctly predicted the searched ribotype when the sourmash search identity cutoff was below 77.7%. Applying a distance cutoff to individual matches comprising concordance or non-concordance to the 4/5 classifier decreased some incorrect calls, identifying them instead as unknowns (Table 3). Sensitivity excluding test genomes with ribotypes not present 4 times or more in the search database was 92.4% (872/944). Of the 78 test ribotypes that could not be determined with this rule, 30 were due to lack of sufficient presence in the database. The 50 incorrect predictions were from 33 non-redundant searched and matched ribotype pairs, of which 26 shared an ST. For the 5 most common ribotypes, RT002 and RT015 were always correctly predicted (both 65/65), RT027 was almost universally correctly predicted (196/197 with one deemed undeterminable), with high rates also for RT078 (86/90; 2 incorrect and 2 undeterminable). RT014 was predicted correctly in 78/87 searches with 5 incorrect and 4 undeterminable predictions.

**Table 3.** Performance of ribotype prediction for 1000 genomes. Taking the 5 closest matches to a database of genome signatures (sourmash search) or full genomes (alignment) and predicting the searched genomes ribotype based on 5/5 or 4/5 concordance. Results would be reported as inconclusive when they lacked concordance, correct when the concordant ribotypes matched the searched and incorrect when they didn’t. A percent cutoff was further applied below which no individual matches had the same ribotype as the searched genome signature

| **Rule** | **Correctly determined** | **Incorrectly determined** | **Determined inconclusive** |
| --- | --- | --- | --- |
| **Sourmash search closest 5/5** | 780 | 20 | 200 |
| **Sourmash search closest 4/5** | 872 | 50 | 78 |
| **Sourmash search closest 5/5 and % identity cutoff** | 780 | 20 | 200 |
| **Sourmash search closest 4/5 and % identity cutoff** | 872 | 45 | 83 |
| **SNP distance closest 5/5** | 792 | 20 | 188 |
| **SNP distance closest 4/5** | 863 | 31 | 106 |

We also compared predictions from core SNP distances to evaluate the ground truth of this distance-based approach using both rules, as opposed to the ability of sourmash search to correctly identify the most similar genomes. Requiring 4/5 closest genomes in terms of SNP distance to match the test genome’s ribotype resulted in 863 (86%) correct ribotype predictions, 106 unknown (11%) and 31 incorrect (3%) (Table 3). The number of correct predictions was nine fewer than from sourmash search but with 19 fewer incorrect predictions.

## Discussion

The use of genomic epidemiology for pathogen surveillance depends on the ability to compare potentially large numbers of genomes that may also be diverse. Hashing k-mers and randomly subsampling is fast and scalable making it possible to compare large numbers of genomes. It has previously been shown that serovars of *Salmonella* can be differentiated with mash with high accuracy(36). Building on these findings we investigated whether even more similar *C. difficile* genomes could be identified with k-mer hash set based approaches (full Jaccard distance) and k-mer hash subsampling (MinHash). In real genome data both full k-mer sets and k-mer subsamples comparisons were partially successful. We investigated with simulated genomes, replicate sequences and by correlation with real genome features the possible main performance factors.

Using a diverse set of *C. difficile* genomes, we demonstrate k-mer based approaches can be used to reduce the search space for subsequent comparisons by identifying potentially closely related genomes. The performance observed for highly similar genomes required using genome assembles rather than raw reads, filtering assemblies to avoid contamination, removal of excessively fragmented assemblies and removal of small contigs (<1000bp). Removing small contigs prevented some outlying results where genomes appeared more distantly related than they were by sourmash distance. Many genome pairs with small SNP distances also reported many whole gene differences that were correlated with increased sourmash distance. Similarly, we observed up to several hundred reported gene differences in sequences generated from the same isolate or same pool of extracted DNA, raising questions as to how many reported gene differences are relevant to transmission analysis versus artefactual from the sequencing process. Simulations of genome pairs without accessory genes suggested that sequencing errors could be a major driver of poor concordance between SNP and sourmash distance.

In a diverse dataset, clustering genomes by MinHash could rapidly exclude the majority of dissimilar genome pairs from further alignment and fine-scaled analysis. The genomes clustered by MinHash were comprised of more similar pairs, comparable with genomes clustered by fractional typing schemes such as ribotyping. While ribotyping is not readily possible *in silico*, 7 gene MLST is, including from raw sequencing reads. MLST carries the disadvantage of incomplete schemes and un-typeable genomes but the benefit of known inconclusive results. A MinHash threshold for clustering could be suggested from our results for *C. difficile* diverse lineage exclusions, but the possibility of false negatives (even greater than observed in this study) cannot be excluded. However, MinHash clustering of similar genomes and further alignment could be conducted and rapidly correctly identify outbreaks while alignment-based investigation of the full sample of genomes is ongoing.

We also explored the possibility of a rapid distance-based ribotype prediction method. Subsampling k-mers, using sourmash search and a match consensus rule from a database (sourmash index) of genomes with known ribotypes successfully identified the correct ribotype for most searched genomes. The performance of this method was comparable to using core genome SNP distances, such that the speedup from MinHash did not introduce error. It’s possible that with greater numbers of ribotypes represented, each by a greater diversity of genomes from within that lineage performance of this method could be improved. Direct *in silico* prediction of all known ribotypes however (as with MLST) remains an unsolved challenge, that may become more tractable as read lengths increase sufficiently to allow more complete genome reconstruction.

In conclusion, k-mer based approaches to comparing *C. difficile* assembled genomes at scale offered modest performance. In a genomic surveillance context where hundreds or thousands of genomes for comparison are becoming routine, it does provide the opportunity to computationally inexpensively and rapidly subset genomes for alignment and outbreak detection while full, fine-scaled investigations are ongoing.

## Author statements

### Authors and contributors

Conceptualisation: MPM, MHW, ASW, DWE

Data curation: MPM, DWE

Formal analysis: MPM, DWE

Visualisation: MPM

Writing – original draft: MPM

Writing – review & editing: MPM, MHW, ASW, DWE

Supervision: MHW, ASW, DWE

### Ethical approval

Previously published and available data were analysed, no new data were collected as part of the study.

