## Supplemental Figures 1-7 for "K-mer based prediction of *Clostridioides difficile* relatedness and ribotypes"

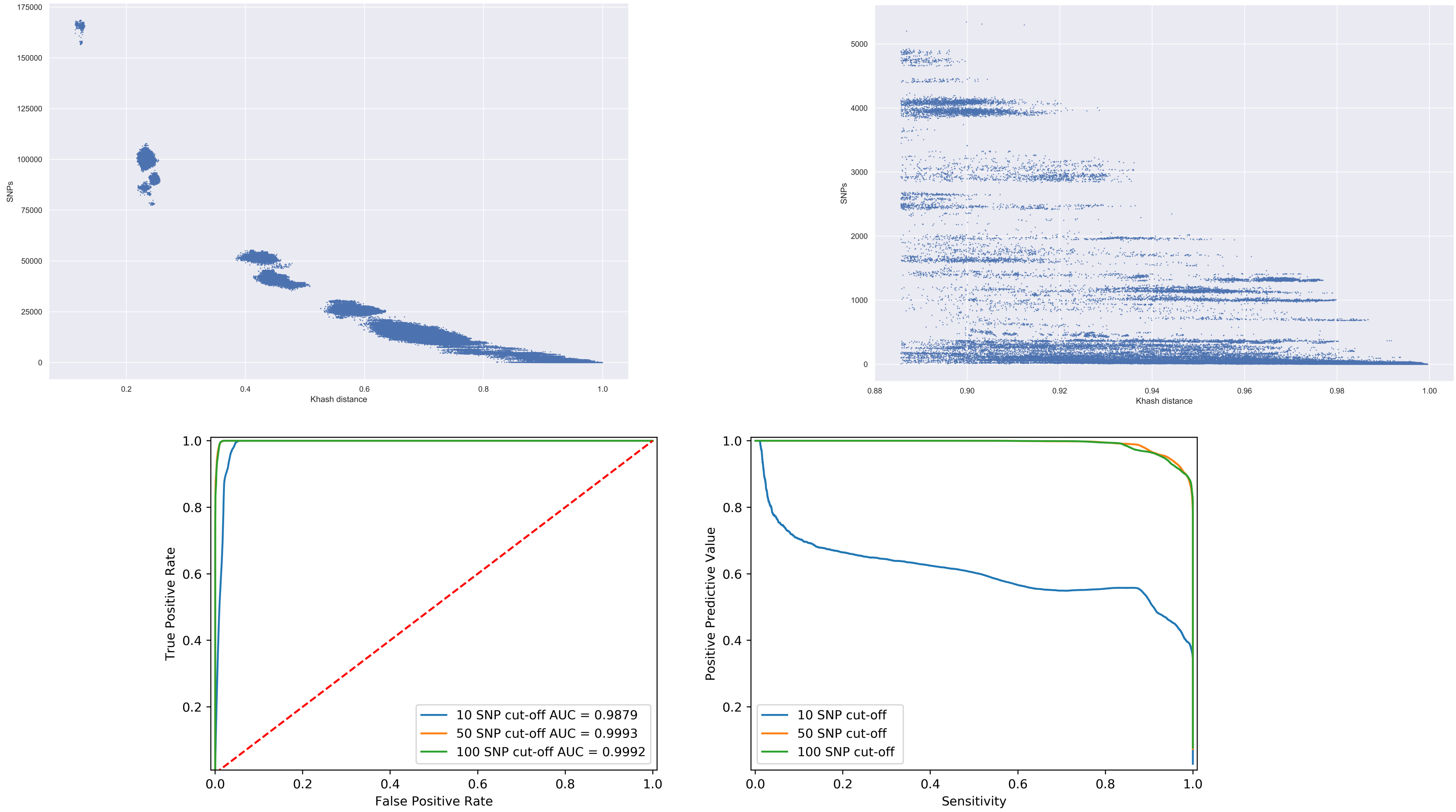


**Supplementary Figure 1.** Scatterplot of SNPs vs full k-mer hash Jaccard distance (khash distance) of 1905 *Clostridioides difficile* genomes comparisons (n = 1,813,560 pairs) (upper left) and the same for those ≥ 0.885652 Khash distance (100% sensitivity for ≤10 SNPs ) (upper right). Performance of sourmash thresholds predicting pairs with ≤10 SNPs (with ≤50 and ≤100 displayed for reference) displayed by receiver operator curves (ROC) with area under curve (AUC) values displayed (lower left) and positive predictive value/sensitivity curves (precision/recall). Colours randomly assigned

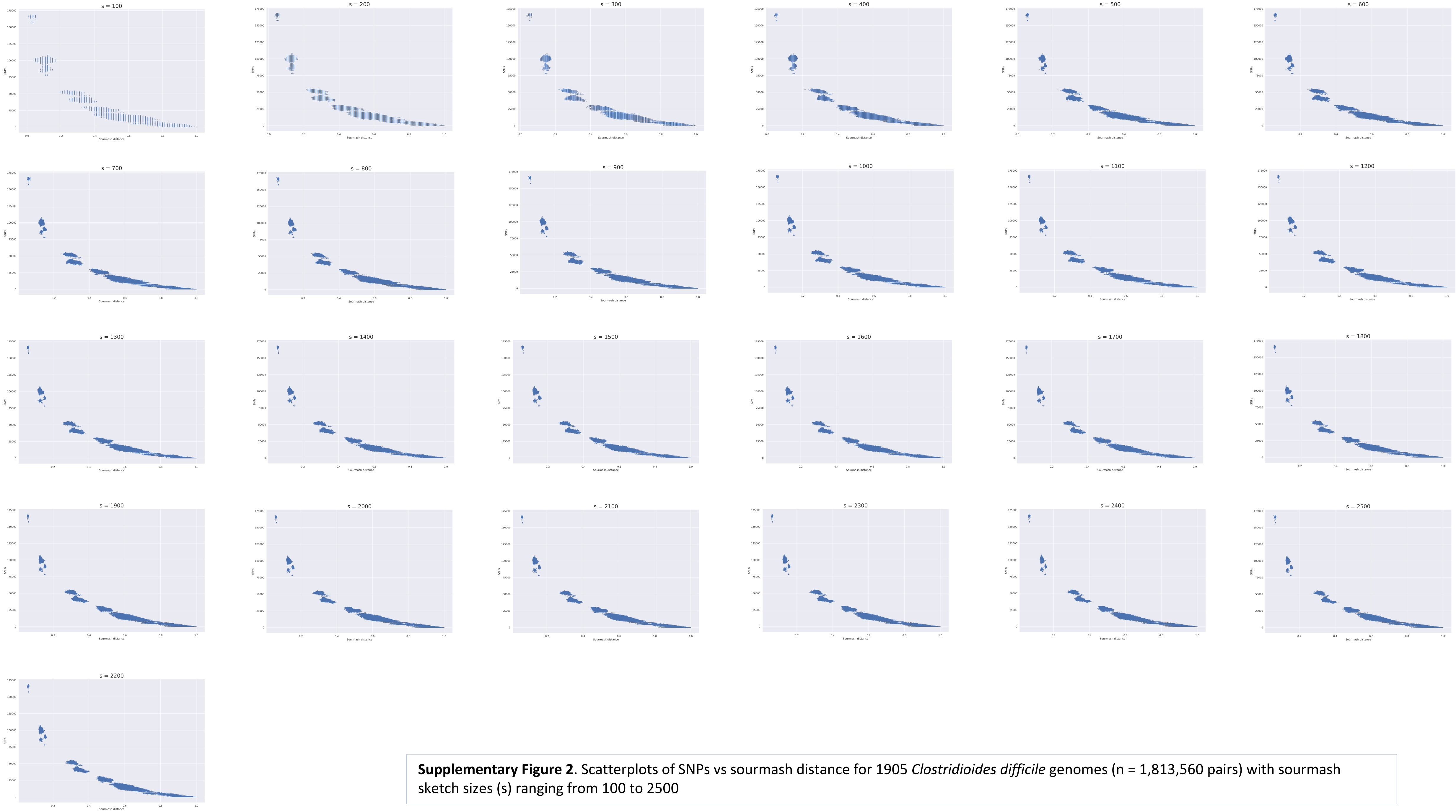


**Supplementary Figure 2**. Scatterplots of SNPs vs sourmash distance for 1905 *Clostridioides difficile* genomes (n = 1,813,560 pairs) with sourmash sketch sizes (s) ranging from 100 to 2500

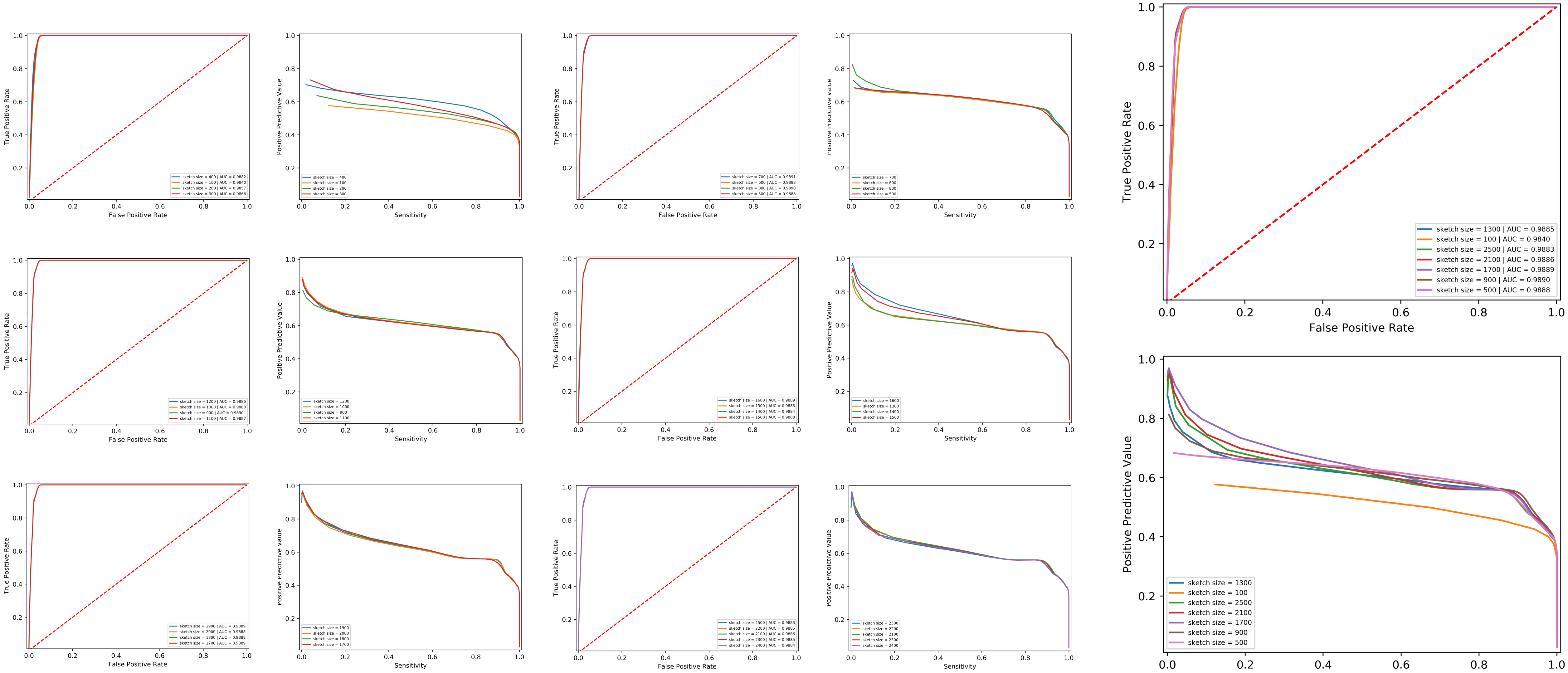


**Supplementary Figure 3**. Performance of sourmash thresholds predicting pairs with ≤10 SNPs displayed by receiver operator curves (ROC) with area under curve (AUC) values displayed and positive predictive value/sensitivity curves (precision/recall). Sourmash sketch sizes (s) 100 to 2500 are displayed. A wide range were sampled and plotted together for reference (upper and lower right). Colours were randomly assigned

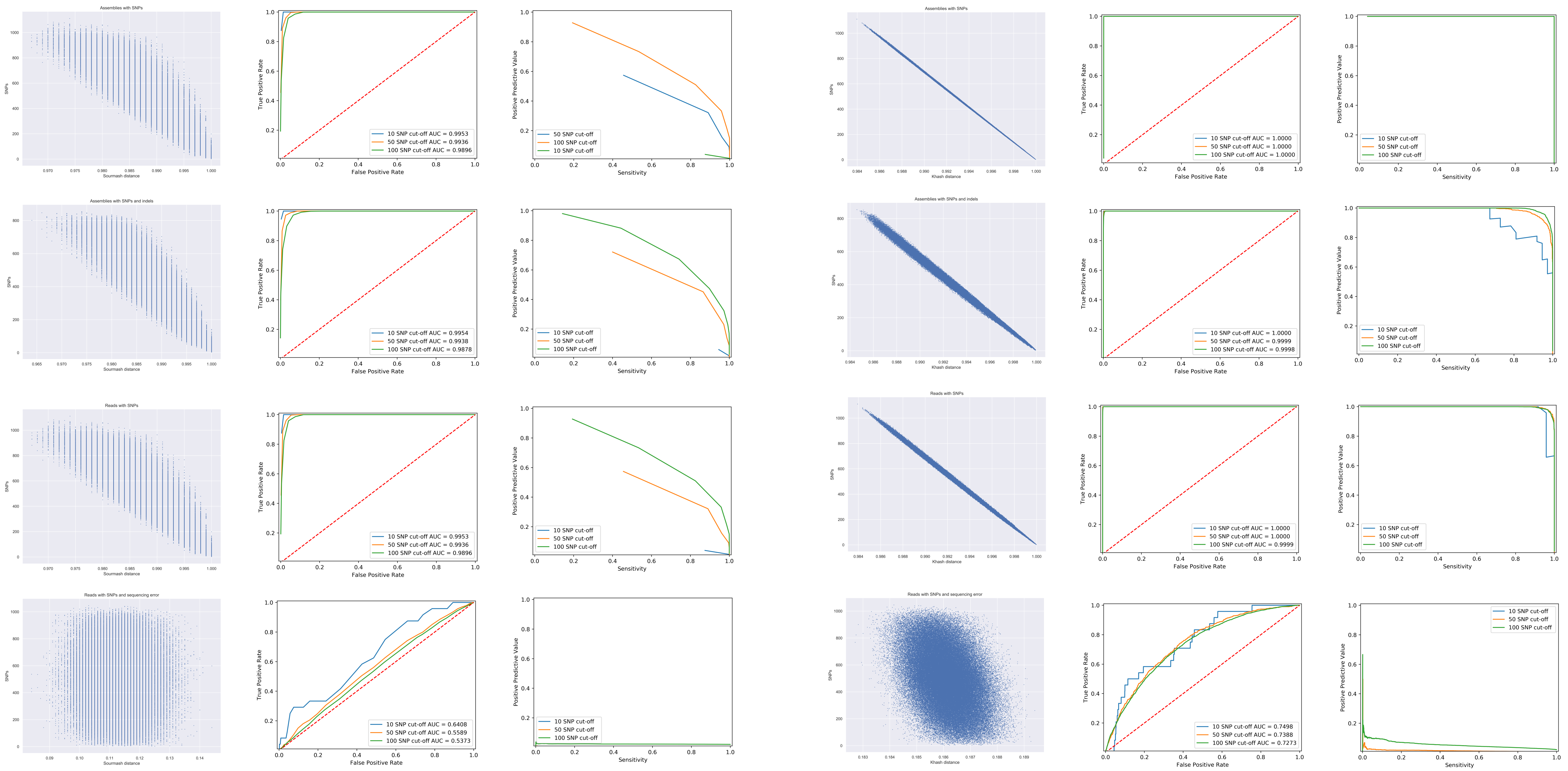

**Supplementary Figure 4.** Performance and distribution of sourmash distances generated from simulated reads or assemblies (n = 500 genomes, 124,750 pairs) and dashing khash full k-mer Jaccard distances for comparison. Performance displayed by receiver operator curves (ROC) with area under curve (AUC) values displayed and positive predictive value/sensitivity curves (precision/recall). On each line performance of sourmash distances predicting <=10 SNPs (<=50, <=100 for comparison) from simulated assembly pair comparisons with SNPs only (left 3 tiles) and full dashing khash distance performance predicting ≤ 10 SNPs (with ≤50 and ≤100 displayed for reference) (right 3 tiles). Lines reading upper to lower are simulated assemblies with SNPs only, simulated assemblies with SNPs and indels, sequencing reads with SNPs only and sequencing reads with SNPs and errors. Colours were randomly assigned

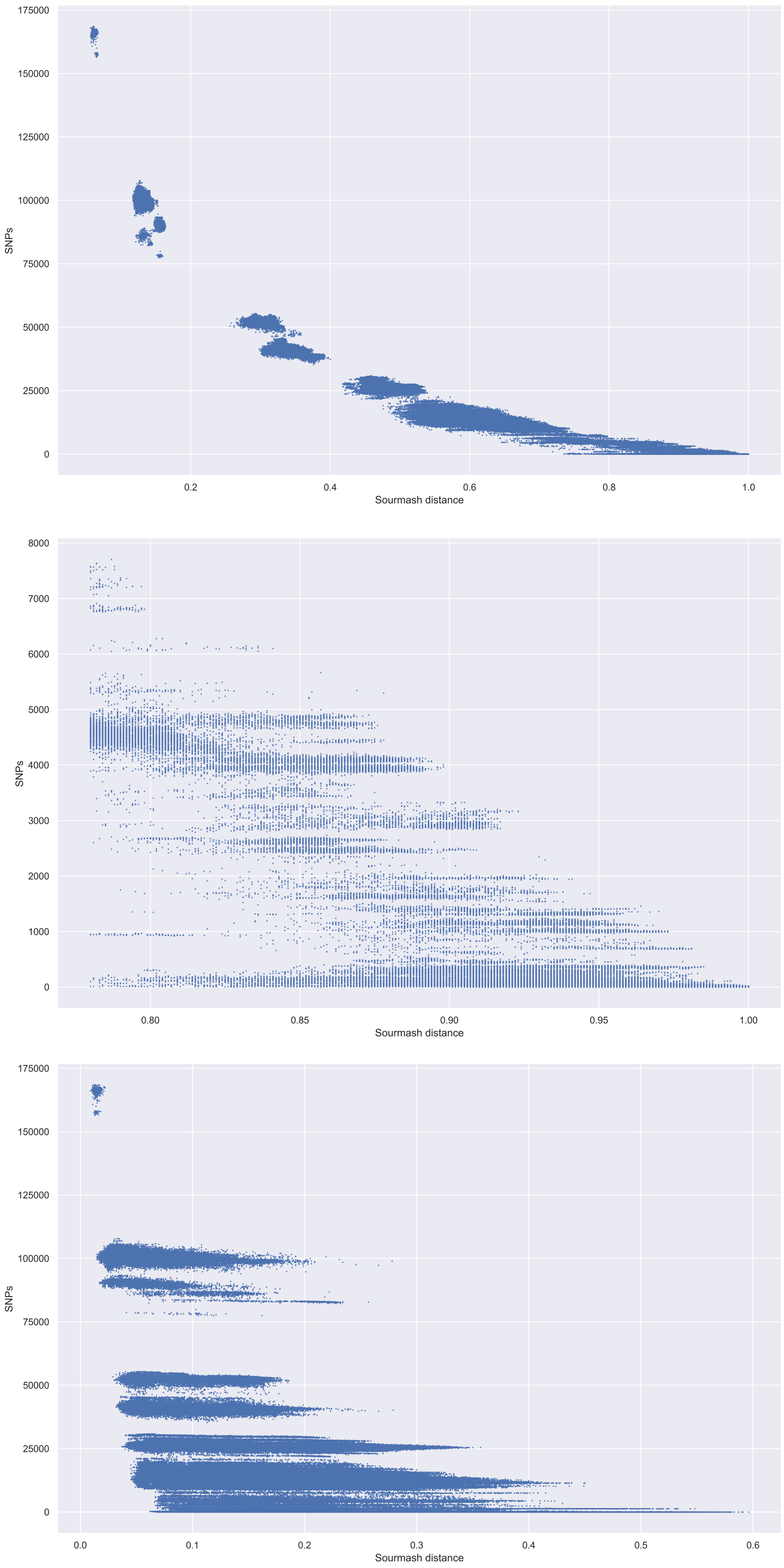


**Supplementary Figure 5**. Scatterplot of genome pair SNPs vs sourmash distance for 1905 *Clostridioides difficile* genomes comparisons (n = 1,813,560 pairs) with low abundance k-mers excluded prior to sourmash sketch generation (upper) and plotted for those ≥0.780 sourmash distance (100% sensitivity for ≤10 SNPs )(middle). SNPs vs sourmash distance for 1905 *Clostridioides difficile* genomes comparisons with prior exclusion of low abundance k-mers (lower)

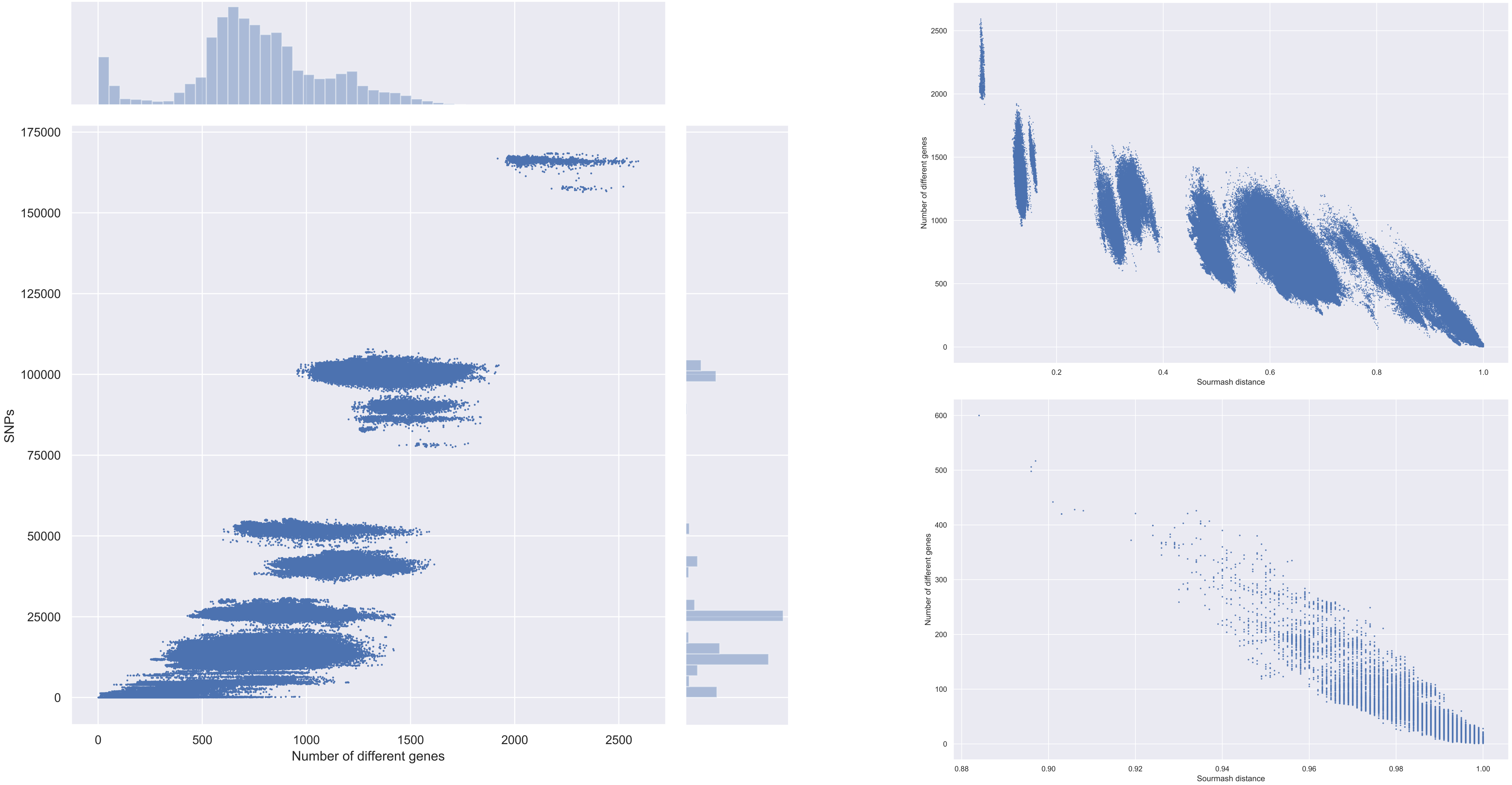


**Supplementary Figure 6**. The relationship between sourmash distance, SNPs and number of different genes of 1905 *Clostridioides difficile* genomes comparisons (n = 1,813,560 pairs). Scatterplot of SNPs vs number of different genes with respective histograms on the margins (left) number of different genes vs sourmash distance (upper right) and number of different genes vs sourmash distance for pairs with ≤10 SNPs (lower right)

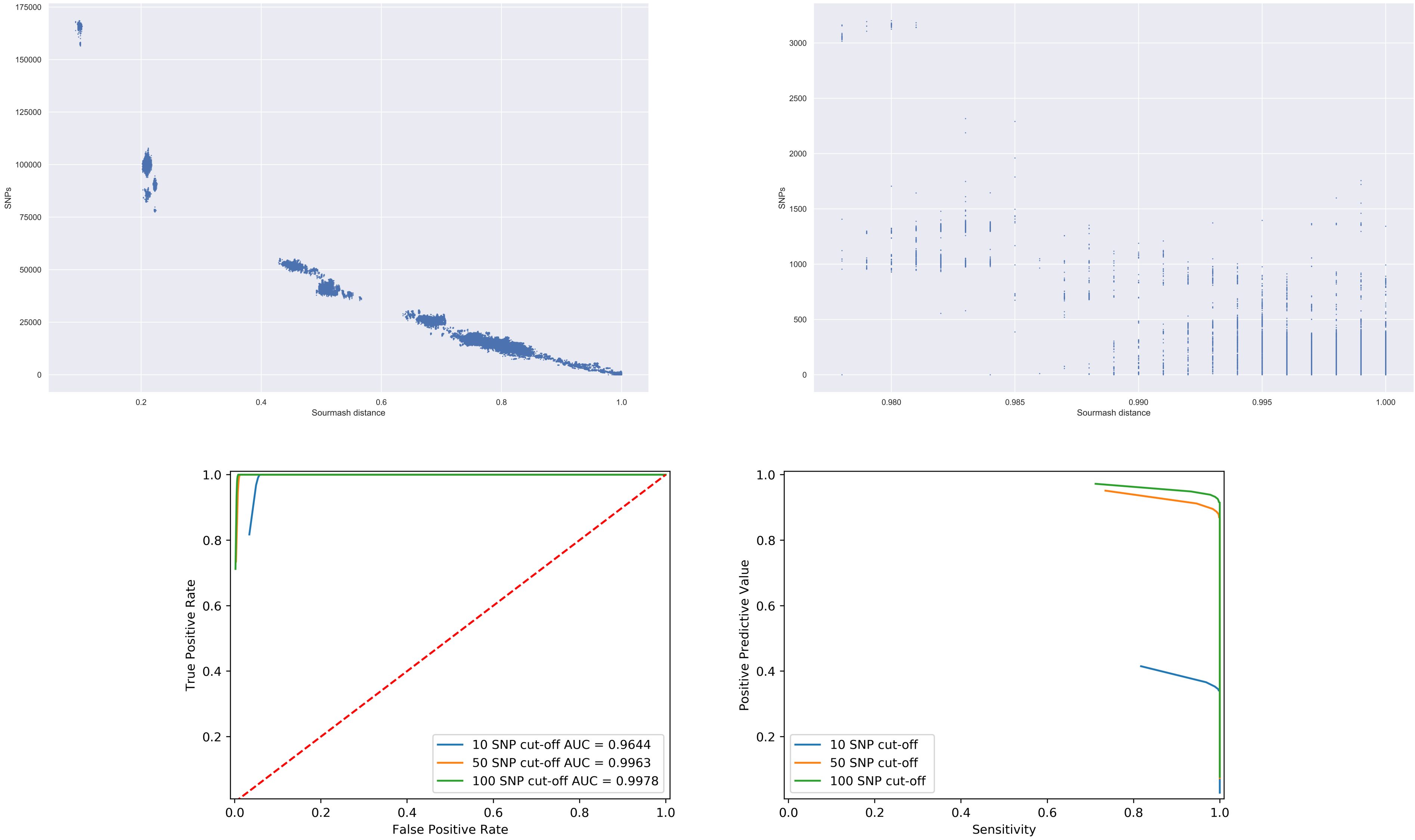


**Supplementary Figure 7**. Scatterplot of genome pair SNPs vs sourmash distance for 1905 *Clostridioides difficile* genomes comparisons (n = 1,813,560 pairs) with k-mers generated from core genes (upper left) and plotted for those ≥0.978 sourmash distance (100% sensitivity for ≤10 SNPs ) (upper right). Performance predicting ≤10 SNPs for all sourmash thresholds receiver operator curve with area under the curve (AUC) values displayed (lower left) and positive predictive value vs sensitivity (precision/recall) curve (lower right). Colours randomly assigned
